# Optimised use of Oxford Nanopore Flowcells for Hybrid Assemblies

**DOI:** 10.1101/2020.03.05.979278

**Authors:** Samuel Lipworth, Hayleah Pickford, Nicholas Sanderson, Kevin K Chau, James Kavanagh, Leanne Barker, Alison Vaughan, Jeremy Swann, Monique Andersson, Katie Jeffery, Marcus Morgan, Timothy EA Peto, Derrick W Crook, Nicole Stoesser, A Sarah Walker

## Abstract

Hybrid assemblies are highly valuable for studies of *Enterobacteriaceae* due to their ability to fully resolve the structure of mobile genetic elements, such as plasmids, which are involved in the carriage of clinically important genes (e.g. those involved in AMR/virulence). The widespread application of this technique is currently primarily limited by cost. Recent data has suggested that non-inferior, and even superior, hybrid assemblies can be produced using a fraction of the total output from a multiplexed nanopore (Oxford Nanopore Technologies [ONT]) flowcell run. In this study we sought to determine the optimal minimal running time for flowcells when acquiring reads for hybrid assembly. We then evaluated whether the ONT wash kit might allow users to exploit shorter running times by sequencing multiple libraries per flowcell. After 24 hours of sequencing, most chromosomes and plasmids had circularised and there was no benefit associated with longer running times. Quality was similar at 12 hours suggesting shorter running times are likely to be acceptable for certain applications (e.g. plasmid genomics). The ONT wash kit was highly effective in removing DNA between libraries. Contamination between libraries did not appear to affect subsequent hybrid assemblies, even when the same barcodes were used successively on a single flowcell. Utilising shorter run-times in combination with between-library nuclease washes allows at least 36 *Enterobacteriaceae* isolates to be sequenced per flowcell, significantly reducing the per isolate sequencing cost. Ultimately this will facilitate large-scale studies utilising hybrid assembly advancing our understanding of the genomics of key human pathogens.

**Data Summary:**

1. Raw sequencing data is available via NCBI under project accession number PRJNA604975. Sample accession numbers are provided in table S1.
2. Assemblies are available via Figshare https://doi.org/10.6084/m9.figshare.11816532.v1.

**Impact Statement:** Most existing sequencing data has been acquired from short-read platforms (eg. Illumina). For some species of bacteria, clinically important genes, such as those involved in antibiotic resistance and/or virulence, are carried on plasmids. Whilst Illumina sequencing is highly accurate, it is generally unable to resolve complete genomic structures due to repetitive regions. Hybrid assembly uses long reads to scaffold together short-read contigs, maximising the benefits of both technologies. A major limiting factor to using hybrid assemblies at scale is the cost of sequencing the same isolate with two different technologies. Here we show that high-quality hybrid assemblies can be created for most isolates using significantly shorter run-times than are currently standard. We demonstrate that a simple washing step allows several libraries to be run on the same flowcell, facilitating the ability to take advantage of shorter running times. Adding nuclease means that contamination between libraries is minimal and has no significant effect on the quality of subsequent hybrid assemblies. This approach reduces the cost of acquiring long reads by >30%, paving the way for large-scale studies utilising hybrid assemblies which will likely significantly enhance our understanding of the genomics of important human pathogens.

## Introduction

Ideally, a single sequencing technology would provide both highly accurate and structurally complete genomes. The rapid acceleration in whole genome sequencing over the past decade has been driven primarily by short-read technologies (e.g. Illumina). The 100-300bp reads generated are generally highly accurate and low cost, and the tools for their analysis are now relatively mature. However, the inability to resolve long genomic repeats using short reads is a significant limiting factor. In *Enterobacteriaceae*, clinically important genes, such as those involved in antimicrobial resistance (AMR) and virulence, are commonly carried on plasmids and other mobile genetic elements (MGEs) [1]. It is generally impossible to delineate the structure of these using short read-data alone [2].

Long-read sequencing platforms such as Oxford Nanopore Technologies (ONT) or Pacific Biosciences (PacBio) can produce reads which are thousands or tens of thousands (and even hundreds of thousands) of bases long. This greatly aids *de novo* assembly because these reads span long genomic repeats. Particularly in the case of ONT however, longer reads are still currently associated with a higher error rate, which may be problematic for some applications (e.g. transmission inference). Improvements in laboratory and bioinformatic methods to enable sequencing using only long-reads are emerging at a rapid pace. Significant limitations remain however, and there has been little evaluation on real world data [3]. Hybrid approaches combine the low error rate of Illumina reads with the structural resolution of ONT/PacBio, maximizing the strengths of both technologies [4] and the widely used Unicycler tool [5] offers an automated and easy-to-use pipeline for this. Large-scale studies utilising hybrid assemblies would likely provide valuable new insights into the biology of MGEs in *Enterobacteriaceae*; however the significant associated cost currently limits the widespread application of this technique.

Recent research has suggested that random subsampling of ONT reads can improve hybrid assemblies [6], raising the possibility that significantly shorter sequencing times may be suitable where long reads are being created for the purpose of hybrid assembly. Producing sufficient reads to complete hybrid assemblies for one library of isolate-extracts may only require a small proportion of the potential useful sequencing time of a flowcell. In theory therefore, it should be possible to sequence multiple libraries on each flowcell, thereby reducing the per-isolate cost. The major obstacle to this is the need to eliminate contamination between libraries sequenced sequentially on the same flowcell. ONT have recently released a version 3 wash-kit with the addition of nuclease. The company quotes between-library contamination as being around 0.1% [7]; however to our knowledge this has not been independently verified.

This study therefore evaluated whether the ONT wash-kit could enable successful re-use of flowcells to increase the number of hybrid assemblies per flowcell for isolates with existing Illumina short-read data. In doing so we investigated: i) whether sequencing run times could be shortened without affecting assembly quality; and ii) whether between-library contamination from reusing flowcells with the new wash kit occurs and can be mitigated. Whilst we primarily focussed on hybrid assembly, we also compared hybrid to long-read only assemblies to assess whether short-read sequencing remains necessary to produce complete and accurate assemblies. Based on these evaluations we propose a rapid and simple workflow which potentially reduces the consumables cost of ONT sequencing by at least 20% with no apparent impact on assembly accuracy.

## Methods

### Isolate preparation, DNA extraction and sequencing

46 isolates were selected for sequencing of which 45 were cultured from bacteraemic patients presenting to Oxford University Hospitals NHS Foundation Trust between 2008-2018, and one was the MGH78578 *Klebsiella pneumoniae* reference (table S1). Pure isolate-cultures were stored at −80°C in 10% glycerol. Sub-cultures of isolate stocks were grown on Columbia blood agar overnight at 37°C. DNA for Illumina sequencing was extracted using the QuickGene DNA extraction kit (Autogen, MA, USA) as per the manufacturer’s instructions with the addition of a mechanical lysis step (FastPrep, MP Biomedicals, CA, USA; 6m/s for 40 secs). Short-read sequencing was performed using an Illumina HiSeq 4000 instrument as previously described[6].

DNA extractions for Illumina and ONT sequencing were prepared from separate sub-cultures. For Nanopore sequencing, DNA from isolates for library 1 (table S1) was extracted using the EasyMag system (bioMerieux). A 10μl loop was used to inoculate 500μl of autoclaved phosphate buffered solution and 100μl of this was transferred to the easyMAG vessel which was then run using the manufacturer’s generic short protocol and a final elution volume of 25μl. For all other extractions for nanopore sequencing, the Qiagen Genomic tip 100/G kit (Qiagen) was used according to the manufacturer’s protocol. DNA concentration was quantified using the Qubit 2.0 instrument (Life Technologies).

DNA extracts were multiplexed as 10 (library 1) or 12 (all other libraries) samples per flowcell using the ONT Rapid Barcoding Kit (SQK-RBK004) according to the manufacturer’s protocol. Sequencing was performed for 48 hours (library 1) and 24 hours for all other libraries on a GridION using version FLO-MIN106 R9.4 flowcells. Flowcells were washed using the ONT Flowcell Wash Kit (EXP-WSH003) and bias voltages adjusted between runs according to the manufacturer’s recommendations. One isolate on library 1 was excluded from all further analysis because the long- and short-read assemblies produced a different species identification, strongly suggesting a laboratory error.

### Read pre-processing and assembly

We compared several filtering and demultiplexing approaches, particularly to try to reclaim ‘unclassified reads’ which might be important when using shorter sequencing times. Overall using Guppy v3.1.5 (https://community.nanoporetech.com) for base-calling and demultiplexing followed by Deepbinner [8] (v0.2.0) to re-assign reads binned as ‘unclassified’ by Guppy produced the most complete assemblies. We therefore adopted this approach for the rest of the analysis (see supplement for details). Quality of ONT reads was assessed by kmer identity compared to Illumina reads using Filtlong [9]. Unicycler v0.4.8-beta was used to create hybrid assemblies utilising both the long- and short-read data. We assessed both Unicycler’s bold and normal ‘--mode’ options (supplement), and elected to use the bold mode results for analysis due to the fact it produced more complete assemblies and a structurally accurate assembly of the MGH78578 reference.

Long-read only assembly was performed using Flye (version 2.6) with the --plasmids option [10]. All assembly graphs were visualised using Bandage [11], which was also used to perform Blastn searches. Isolates (n=3) with <5x estimated genome coverage were excluded from the long read vs hybrid assembly comparison. All computation was performed on the Oxford University Biomedical Research Computing cluster with eight threads used for each assembly. Deepbinner was run on a cluster of NVIDIA GeForce GTX 1080 Ti GPUs.

### Assembly comparison

We compared assemblies created under different conditions using various different metrics:

- Completeness – the number of plasmids/chromosomes in each assembly marked as being circular by Unicycler.
- ALE – assembly likelihood estimator which estimates the likelihood of hybrid assemblies created using the same Illumina short read sequencing data [12]. Short reads were mapped to hybrid assemblies using minimap2 [13].
- DNADiff – whole genome alignment with calculations of gSNP and gIndel differences between assemblies [14]. gSNPs and gIndels represent high confidence SNPs and indels bounded between at least 20 exact nucleotide matches on both sides.

The relationship between the number of long-read bases and completeness (assessed as all structures marked as being circular by Unicycler) was estimated using a Wilcoxon rank sum test in R version 3.6. Minimap2 was used to map contigs from long-read to hybrid assemblies. ML plasmids [15] was used as a further arbitrator of the chromosomal/plasmid origin of sequences. Human reads were detected using Centrifuge [16] as part of the Crumpit [17] pipeline. Simulations of shorter sequencing times were performed by selecting reads from fastq files produced between the beginning of the run and the simulated endpoint using a python script (available at https://github.com/samlipworth/ONT-wash-hybrid).

### Phases of laboratory evaluation

Three laboratory phases were performed (figure 1):

**Figure 1:**
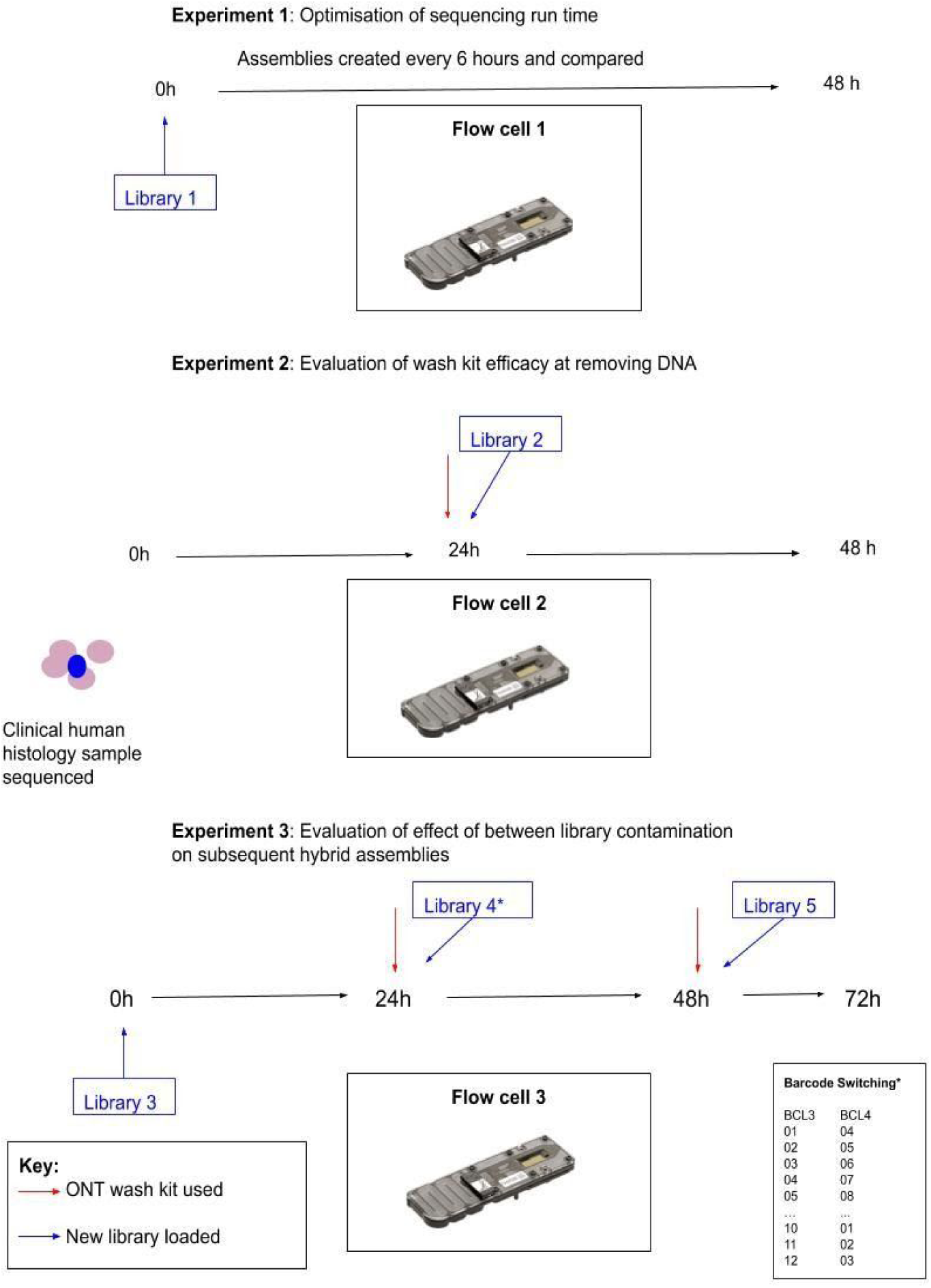
Schematic representation of experiments performed, flowcells used and libraries sequenced. *The same 12 isolates were sequenced in both libraries 3 and 4 however with different barcodes as shown in the inset table and table 1.

1. **Optimisation of flowcell run time** (Flowcell 1, library 1):
  a. 10 isolates (1 excluded from analysis, see above) sequenced for 48 hours with comparison of assemblies created every 6 hours (i.e. the first assembly used the first 6 hours of data and the second the first 12 hours etc.)
2. **Quantification of between-library contamination after using ONT wash kit** (Flowcell 2, library 2):
  a. Assessed by washing and then reusing a flowcell which had been used to sequence a clinical pathology sample for 24 hours for an unrelated project. As library 2 contained 12 pure culture bacterial samples, no human DNA should have been detected if the wash kit is completely effective.
3. **Evaluation of the effect of between library contamination on subsequent hybrid assemblies** (Flowcell 3, libraries 3-5):
  a. Assessed by first sequencing twelve isolates for 24 hours (library 3) and then washing the flowcell and re-sequencing the same 12 isolates with all barcodes switched (library 4, table 1). We subsequently compared hybrid assemblies created using long-read data from libraries 3 and 4.
  b. We then washed flowcell 3 for a second time and sequenced 12 different isolates for a further 24 hours. We checked for between library contamination by blasting contigs (BLASTn) from short-read to hybrid assemblies.

## Results

### Optimisation of sequencing run-time

We ran the first flowcell with library 1 for 48 hours (multiplexing 10 isolates of which 1 was excluded from analysis due to laboratory error). For the nine evaluable isolates, read length peaked at an N50 of 8774 base pairs (bp) after 7 hours and subsequently decreased to a minimum of 7094 bp at 42 hours. Median read quality score peaked at 5 hours (72, IQR 43-85) and reduced to a minimum of 54 (IQR 23-73) at 38 hours. The rate of bases called for each barcode over time was very unequal (figure S1); at twenty-four hours there was a median output of 447 Mb per barcode (range 131Mb – 863Mb) and at 48 hours there was a median of 552 Mb per barcode (range 158 Mb-1085 Mb).

**Table 1:** Comparison of SNPs and indels detected by DNAdiff between hybrid assemblies of the same isolates sequenced in libraries 3 and 4.

| Isolate Name | Barcode library 4 | Barcode library 3 | SNPs | Indels | % Identity | Species | MLST |
| --- | --- | --- | --- | --- | --- | --- | --- |
| blc-44 | 1 | 10 | 9 | 64 | 99.99 | *E. coli* | 420 |
| blc-45 | 2 | 11 | 27 | 1 | 99.99 | *K. pneumoniae* | 490 |
| blc-46 | 3 | 12 | 36 | 2 | 99.98 | *E. coli* | 372 |
| blc-47 | 4 | 1 | 0 | 0 | 100 | *K. pneumoniae* | 490 |
| blc-48 | 5 | 2 | 0 | 0 | 100 | *K. pneumoniae* | 490 |
| blc-49 | 6 | 3 | 0 | 0 | 99.99 | *K. pneumoniae* | 45 |
| blc-50 | 7 | 4 | 0 | 0 | 100 | *E. coli* | 127 |
| blc-51 | 8 | 5 | 0 | 0 | 100 | *K. pneumoniae* | 15 |
| blc-52 | 9 | 6 | 0 | 1 | 99.99 | *E. coli* | 428 |
| blc-53 | 10 | 7 | 0 | 0 | 100 | *E. coli* | 127 |
| blc-54 | 11 | 8 | 1 | 5 | 99.99 | *E. coli* | 88 |
| blc-55 | 12 | 9 | 3 | 1 | 99.99 | *K. pneumoniae* | 490 |

*E. coli* - *Escherichia coli*, *K. pneumoniae* - *Klebsiella pneumoniae*.

To empirically estimate the optimum run time we compared hybrid assemblies produced at cumulative six-hourly intervals during the 48 hours over which library 1 was sequenced. Maximum circularity was achieved by 24 hours by which point 6/9 assemblies (24/27 contigs) had fully circularised (figure 2); notably there was no further benefit gained from an additional 24 hours of sequencing. By 24 hours, 17/18 plasmids had circularised; one was comprised of a single contig but not marked as circular by unicycler (which was also the case at 48 hours). Comparison of the assembly of the reference strain (MGH78578 – barcode 1) at 12 hours (the only time point at which it completely circularised) to the published sequence revealed the correct number of plasmids (n=5) and a high degree of genetic similarity (1 unaligned base, 99.97% average identity, 64 gSNPs and 31 gIndels).

**Figure 2:**
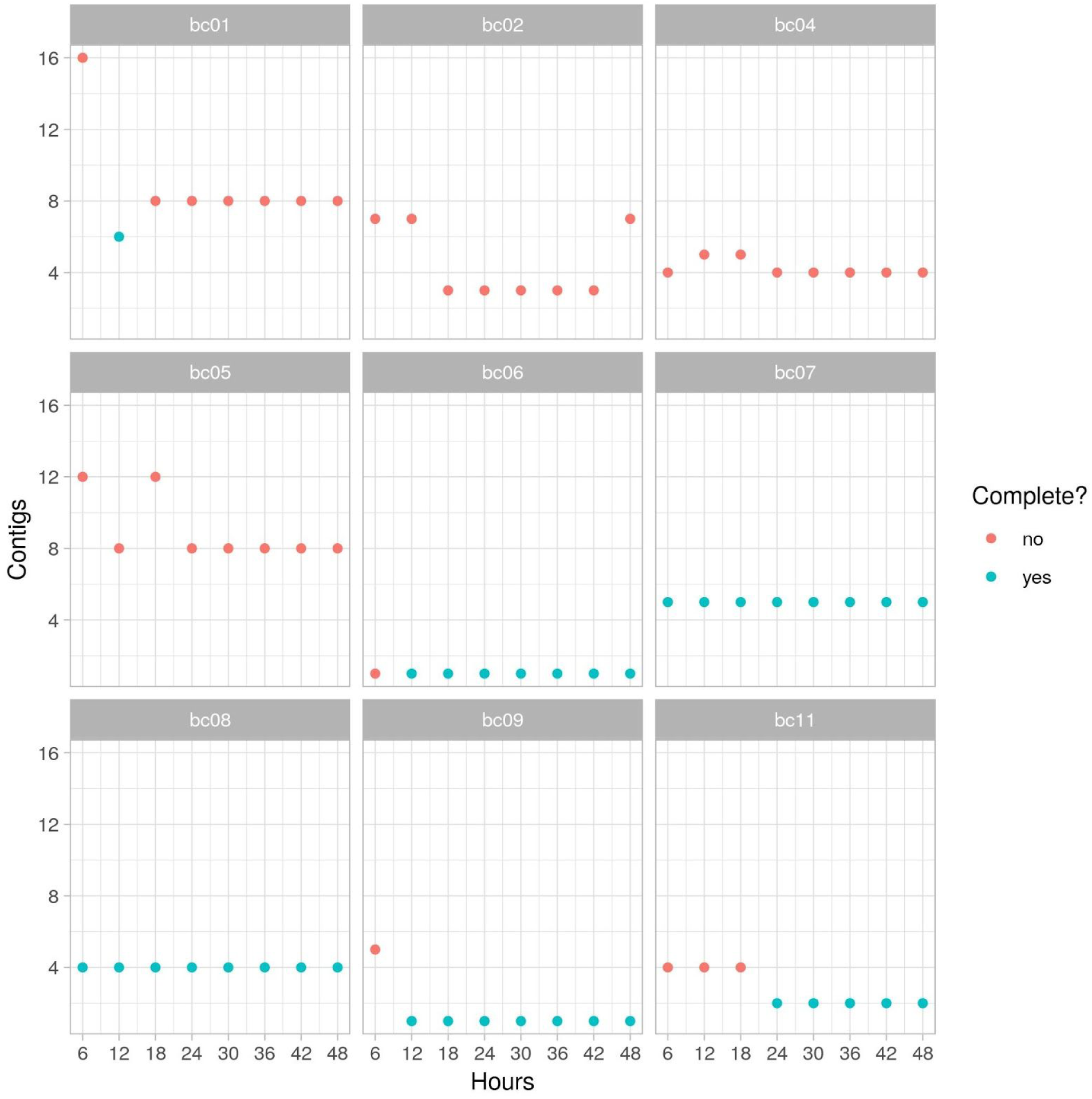
Number of contigs generated by Unicycler for the nine included barcoded (bc) samples in library 1 over time (one isolate was excluded, see methods). Complete assemblies (where the chromosome and all plasmids are formed of single, circularised contigs) are shown in blue.

The three non-complete assemblies (barcodes 02, 04, 05; isolates blc-23, blc-24, blc-25) at 24 hours had complete plasmid structures and relatively simple chromosome graphs (figure S2). There was no relationship between the number of long read bases and probability of hybrid assembly completion at 24 hours (p=0.17), reinforcing the likely futility of longer sequencing times. We also compared the assemblies created at different timepoints using the ALE tool which revealed a similar pattern of results: In two cases (bc04 and bc09, isolates blc-24 and blc-29), a more likely assembly vs that at 48 hours was obtained after 24 hours (figure 3). For the rest of this study we therefore elected to stop all sequencing runs at 24 hours.

**Figure 3:**
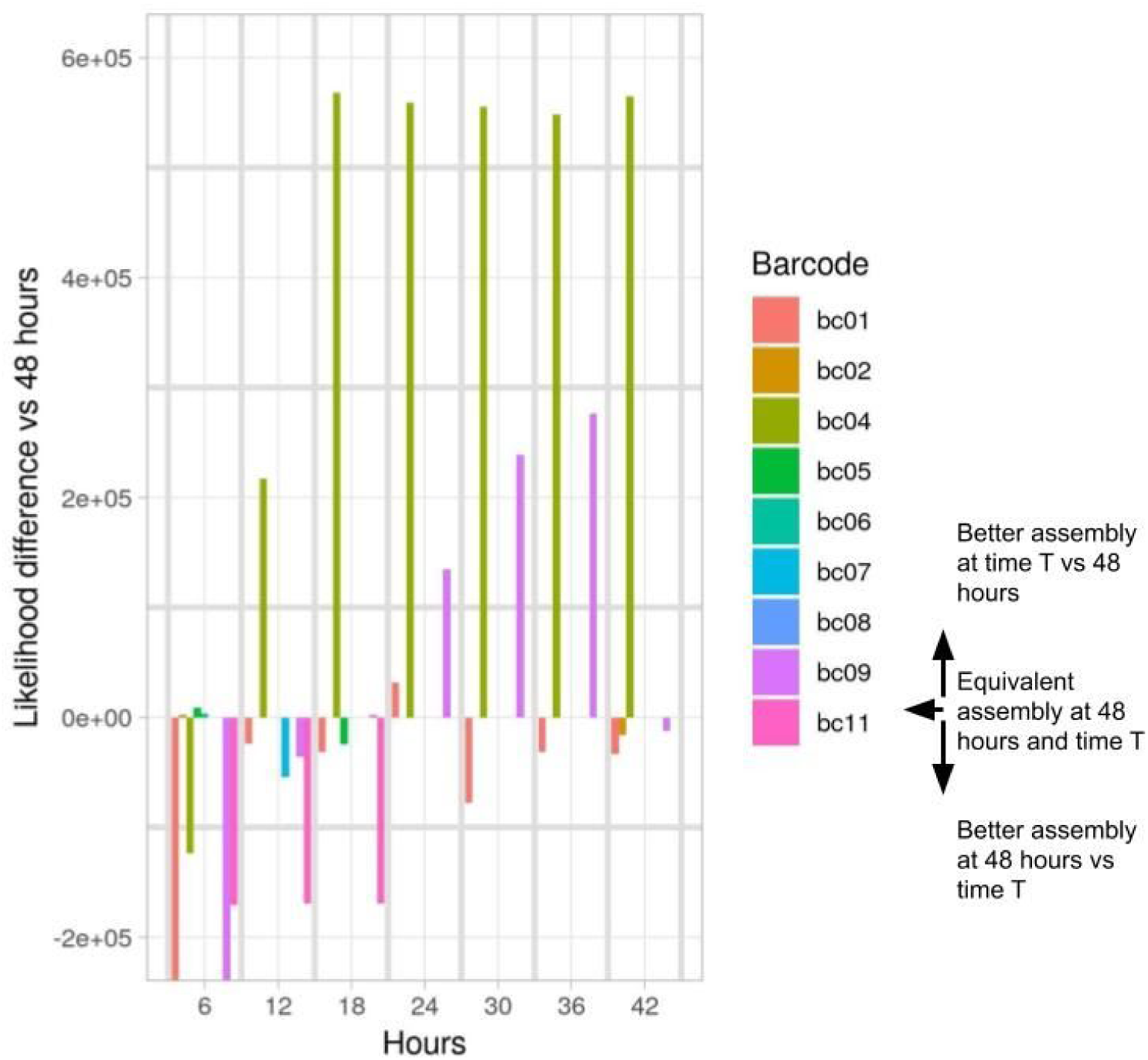
Assembly likelihoods were calculated with the Assembly Likelihood Estimator (ALE) by mapping Illumina reads to hybrid assemblies. Likelihoods were calculated for assemblies of each barcoded isolate created at 6-hour intervals up to 48 hours (x axis). The y-axis denotes likelihood difference between the assembly 48h vs that at time T. A likelihood difference of 0 (and thus no bar-line visible) implies that the assembly at time T is equally as likely as that at T = 48 hours. A positive likelihood difference implies that the assembly at time T was better than at 48 hours.

### Evaluation of wash kit efficacy at removing human DNA

We first attempted to use the ONT wash kit on a flowcell which had previously been used to sequence a human clinical pathology sample for 24 hours. This first 24 hours of sequencing yielded 2,059,966 reads of which 2,028,024 (98.4%) were binned as human by centrifuge. The flowcell was then washed and reloaded with library 2 (bacterial isolates only), which was sequenced for 24 hours. After demultiplexing, 818091 reads (3942 Mb) were obtained of which 147 (0.02%) were binned by centrifuge as being of human origin. The number of human reads was within the range of human reads called by centrifuge for all other flowcells (which had not sequenced any human DNA, table S2), suggesting this number is compatible with background noise from the kit-ome/false-positive binning.

Using this 24-hour-old recycled flowcell to sequence library 2, we acquired complete assemblies for 10/12 genomes. Barcode 1 failed, returning only 3.8×10^6^ bases of data and barcode 10 yielded an assembly of four contigs comprising a chromosome and three plasmids but one of the plasmids was marked as incomplete by Unicycler.

### Reusing flowcells for similar isolates

Given the low contamination observed, we next sought to reuse a flowcell to sequence closely related *Enterobacteriaceae* using a single set of barcodes. After 24 hours, library 3 produced 8/12 fully complete assemblies, following which we re-sequenced the same isolates after changing the barcodes used as shown in table 1, and washing the flowcell between runs. Starting channel availability decreased by about 28% (∼1400 to ∼1000 at the beginning of library 3 vs 4 respectively, figure S3).

Completeness was identical for 11/12 isolates between the runs. There was however a major discrepancy in one sample where a ∼ 868kb region was called as chromosomal in library 3 and a circularised super-plasmid-like component in library 4 (figure 4). As expected, ML plasmids [15] predicted with high confidence (97% probability) this contig was of chromosomal origin. Interestingly this error was fixed after filtering with Filtlong, suggesting it may have arisen from low quality reads.

**Figure 4:**
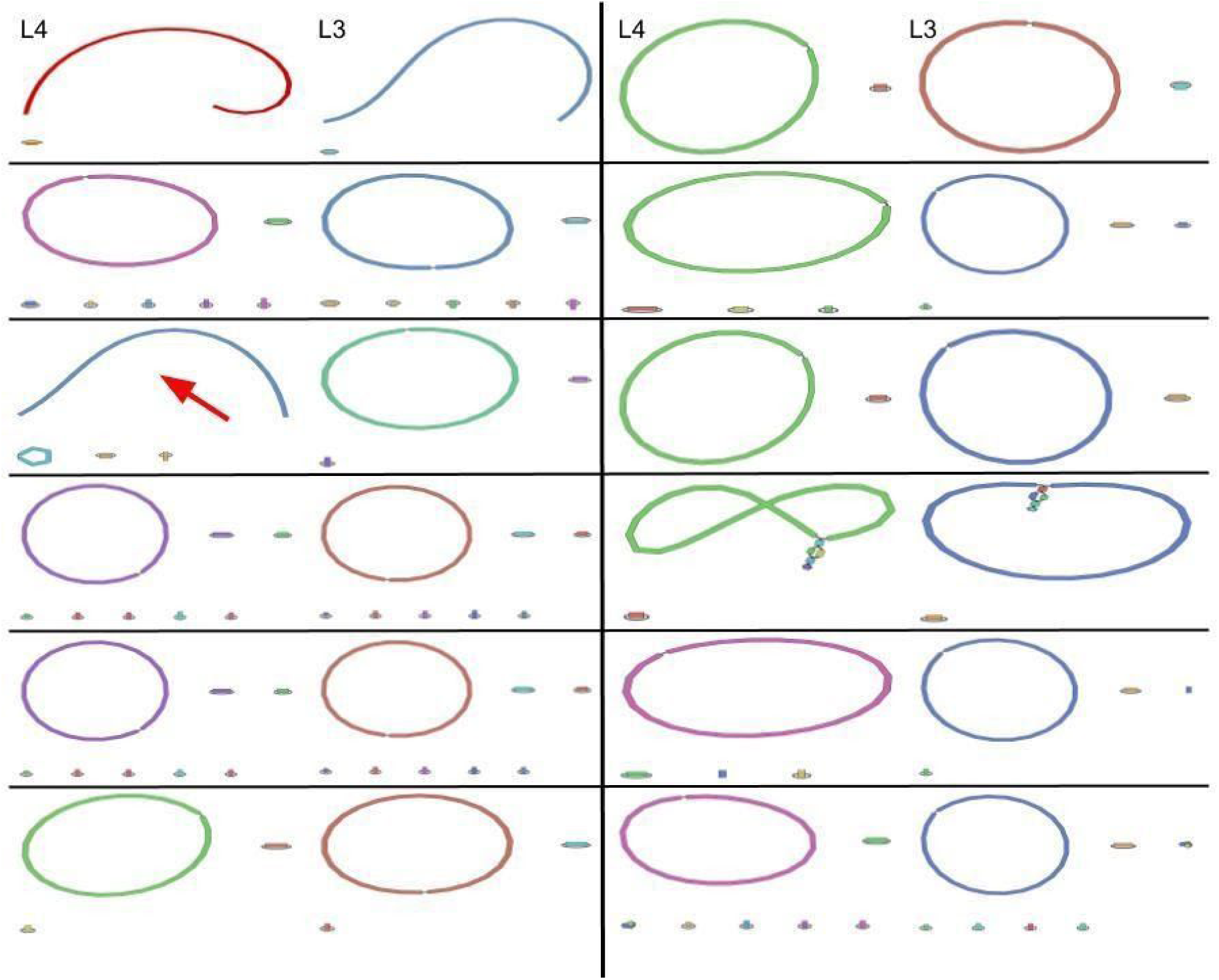
Each box represents the graph of a single isolate sequenced with different barcodes in libraries 3 (L3) and 4 (L4). The left panel contains isolates blc-44-49 and the right L4 isolates blc-50-55 (see table 1). Red arrow denotes the assembly with the major structural difference (blc-46).

For 6/12 isolates (blc-46, blc-48, blc-50, blc-51, blc-53, blc-55,) the ALE score suggested a better assembly for library 3. Comparing hybrid assemblies of the same isolates between libraries (i.e. library 3 vs library 4) using the DNAdiff tool revealed near-identical assemblies (identity >= 99.98%), and low numbers of SNPs and indels in all instances (table 1). The worst performing assembly (blc-46) contained the major structural disagreement discussed above which likely caused the slightly higher number of SNPs between assemblies in this sample. We subsequently reloaded the same flowcell which had been used to sequence libraries 3 and 4 with library 5 (different isolates) and generated a further 7/12 completed assemblies. Blast searches confirmed that all contigs present in these hybrid assemblies were also present in the short-read assemblies; there was no evidence of between-library contamination.

A possible explanation for differences in assemblies between runs might be that, as demonstrated above, read length and quality deteriorates markedly over the course of a single run cycle which may introduce false artificial variation. However use of the ONT wash kit on flowcell 3 between libraries 3 and 4 restored median read quality scores almost to their original values (before = 70 (IQR 10-84), after = 69 (IQR 25-82)). From 18-24 hours of sequencing library 3, median read length was 1204 (IQR 169-4429). After using the wash kit, the next six hours of sequencing of library 4 yielded median read length of 1439 (IQR 292-4554) (figure S4). A similar effect appeared to occur after the flowcell was washed and re-loaded with library 5 although quality scores and read lengths decayed quicker on the third run. However results from library 5 were not directly comparable because they comprised sequencing data from different extractions and isolates.

### Evaluating sequencing run-times using all data

Combining data from libraries 1, 2, 3 and 5 (i.e. excluding library 4 as the isolates were the same as in library 3), we simulated shorter sequencing times by assembling reads produced 3, 6 and 12 hours after the start of the run. At 24 hours, 29/45 isolates had complete assemblies (chromosome and all plasmids circular) compared to 29/45 at 12 hours, 24/45 at 6 hours and 21/45 at 3 hours. The number of incomplete plasmids was similar across all time points (8/150 at 24 hours, 6/150 at 12 hours, 7/150 at 6 hours and 7/150 at 3 hours).

### Comparison with long read assembly

Finally, we compared hybrid assemblies to those generated using only long reads to assess whether generating Illumina reads is still likely to be necessary for future studies. Overall, long-read only assemblies had a high average identity to the reference hybrid assemblies (table 2). When created using data demultiplexed by Guppy alone however, most of the long-read only assemblies contained contigs which did not map to the hybrid assemblies (median number 3, range 0 – 15, median length 3780 bp, range 545 – 19197 bp, median coverage 24, range 4 – 995) (figure S5). This was true both for libraries sequenced on new flowcells and those that had been reused after washing, but not for library 1. Using BLASTn (in Bandage) we were able to identify that some of these likely represented between barcode contamination from isolates sequenced in the same library (figure S6).

**Table 2:** DNAdiff comparisons between Flye long-read only assemblies and hybrid assemblies.

| Library | Isolate Name | Barcode | SNPs | Indels | % reference bases aligned | % Identity |
| --- | --- | --- | --- | --- | --- | --- |
| 1 | blc-22 | bc01 | 3385 | 14679 | 99.95 | 99.55 |
| 1 | blc-23 | bc02 | 251 | 9953 | 100 | 99.76 |
| 1 | blc-24 | bc04 | 3202 | 15949 | 99.98 | 99.58 |
| 1 | blc-25 | bc05 | 3387 | 12262 | 100 | 99.53 |
| 1 | blc-26 | bc06 | 3082 | 11494 | 99.96 | 99.53 |
| 1 | blc-27 | bc07 | 3203 | 11764 | 99.98 | 99.56 |
| 1 | blc-28 | bc08 | 157 | 12793 | 99.97 | 99.61 |
| 1 | blc-29 | bc09 | 3309 | 12036 | 100 | 99.39 |
| 1 | blc-31 | bc11 | 4070 | 13861 | 99.98 | 99.51 |
| 2 | blc-33 | bc02 | 3179 | 15572 | 99.97 | 99.48 |
| 2 | blc-34 | bc03 | 4942 | 16498 | 100 | 99.34 |
| 2 | blc-35 | bc04 | 5488 | 16499 | 99.9 | 99.36 |
| 2 | blc-36 | bc05 | 5458 | 16842 | 100 | 99.35 |
| 2 | blc-37 | bc06 | 5442 | 16499 | 99.98 | 99.36 |
| 2 | blc-38 | bc07 | 5460 | 16559 | 100 | 99.36 |
| 2 | blc-39 | bc08 | 4311 | 20783 | 97.82 | 98.45 |
| 2 | blc-40 | bc09 | 5428 | 16882 | 99.83 | 99.36 |
| 2 | blc-41 | bc10 | 3064 | 15975 | 99.93 | 99.37 |
| 2 | blc-42 | bc11 | 2960 | 15133 | 100 | 99.44 |
| 2 | blc-43 | bc12 | 5434 | 16748 | 100 | 99.35 |
| 3 | blc-47 | bc01 | 4482 | 19204 | 99.98 | 99.31 |
| 3 | blc-48 | bc02 | 4352 | 19444 | 99.99 | 99.3 |
| 3 | blc-49 | bc03 | 3983 | 18206 | 99.98 | 99.3 |
| 3 | blc-50 | bc04 | 2615 | 17537 | 100 | 99.4 |
| 3 | blc-51 | bc05 | 3728 | 19560 | 100 | 99.36 |
| 3 | blc-52 | bc06 | 2645 | 16326 | 100 | 99.4 |
| 3 | blc-53 | bc07 | 2645 | 17567 | 100 | 99.4 |
| 3 | blc-48 | bc08 | 2580 | 17145 | 99.95 | 99.39 |
| 3 | blc-49 | bc09 | 4326 | 19080 | 99.95 | 99.31 |
| 3 | blc-50 | bc10 | 2607 | 15458 | 100 | 99.4 |
| 3 | blc-51 | bc11 | 4406 | 19273 | 99.99 | 99.31 |
| 3 | blc-52 | bc12 | 2555 | 17336 | 100 | 99.4 |
| 5 | blc-56 | bc01 | 4746 | 19881 | 99.89 | 98.96 |
| 5 | blc-57 | bc02 | 4174 | 20993 | 99.46 | 98.56 |
| 5 | blc-58 | bc03 | 4876 | 20019 | 99.92 | 99.01 |
| 5 | blc-59 | bc04 | 4649 | 21014 | 94.41 | 97.93 |
| 5 | blc-61 | bc06 | 3115 | 14943 | 100 | 99.45 |
| 5 | blc-62 | bc07 | 4709 | 15872 | 100 | 99.33 |
| 5 | blc-63 | bc08 | 4680 | 15201 | 99.98 | 99.34 |
| 5 | blc-64 | bc09 | 3051 | 15973 | 100 | 99.39 |
| 5 | blc-65 | bc10 | 4212 | 18101 | 99.36 | 99.22 |
| 5 | blc-66 | bc12 | 4986 | 21189 | 98 | 98.61 |

To try to correct this, we created further assemblies using only reads where both Deepbinner and Guppy agreed on the barcode assignment. Whilst this greatly improved the assemblies and most (but not all) spurious contigs were removed (figure S5), structural differences compared to the hybrid references remained in several assemblies (figure S7). We hypothesised that this might be an issue with rapid barcoding but saw the same signal in data multiplexed with the native barcoding kit in a recent study [6] (figure S8).

## Discussion

In this study we have demonstrated that, for the purposes of creating ONT reads from pure isolates for hybrid assembly, there is unlikely to be benefit in extending sequencing runs beyond 24 hours; indeed, for most assemblies 12 hours is likely to be sufficient. We have also shown that after utilising the ONT flowcell washkit, between library contamination is minimal, and is unlikely to have an important effect on subsequent hybrid assemblies. This appears to be true even when the same barcodes are used for successive libraries. Despite significantly shortened run-times and reusing flowcells, we were able to completely assemble the vast majority of plasmids. This marks a significant milestone for ONT sequencing for the purposes of hybrid assembly and unlocks the potential for large-scale studies of plasmid epidemiology in the near future.

Previous studies have demonstrated successful completion of 12 genomes on a single flowcell; here we have demonstrated this can be increased to at least 22. Based on ONT’s quoted figures of £500 per flowcell and £150 for library preparation and barcoding, the current per sample cost for the long-read sequencing component is £54 (i.e. £500 for flowcell + £150 for library preparation/12). In the most conservative interpretation of this study, we have shown an approximately 33% per sample reduction in cost to £36 (i.e. £500 for flowcell + 2x£150 for library preperation/22), assuming downstream analysis demanded complete circularisation of all contigs. We envisage that for most current use cases however, particularly plasmid genomics, the standard of data produced in the majority of our assemblies would be sufficient to answer the biological questions posed. Even with ultra-short run times of 12 hours (1/6th of the total run-time that is currently standard in our lab and others) we were able to circularise the vast majority of plasmids (and most chromosomes).

If as seems plausible from our data, 12 hours is a viable run-time for most research questions and we assume a useful period of 72 hours per flowcell, then long-read sequencing costs would be further reduced to approximately £19.40 per isolate (i.e. £500 for a flowcell + 6 × £150 for library prep /72 isolates, a ∼ 64% reduction on current costs). This might be limited by the effect of repeated washing of the flowcell and deterioration of pores over time; however even in library 4 in our study (which used a 48 hour old flowcell which had been washed twice), 8/12 chromosomes and 33/36 plasmids were complete at 12 hours. We envisage that after stopping runs at 12 or 24 hours, investigators would be able to carefully select the few isolates which require further sequencing and avoid wasting valuable pore time where complete assemblies have already been acquired. We would caution however that, in this study, increasing run-times did not usually lead to improved assemblies. This is consistent with recent data from a different study in our laboratory which demonstrated that in some cases random sub-sampling of reads can even improve assemblies [6].

Ideally, one would want to use a unique set of barcodes for each library run on a single flowcell. At present however there are only 12 barcodes available in ONT’s rapid barcoding kit which has a substantially easier and less time consuming protocol compared to the Native Barcoding Kit (for which 24 barcodes are available). Nevertheless the number of SNPs and indels between alignments of the isolates sequenced in different libraries on the same flowcell (using the same set of barcodes but reassigned to different isolates) was similar to that seen comparing Illumina/ONT and Illumina/PacBio assemblies of a single isolate [6]. Our assembly of the MGH757878 reference diverged by a similar number of SNPs compared to the published sequence and that in a recent study [6]. To our knowledge there are limited data available on the variation produced by successive cycles of culturing, DNA extraction and sequencing the same isolate using ONT technology and further investigation of this using reference sequences seems warranted. Based on our data, using the same barcodes for consecutive libraries on the same flowcell is likely to be acceptable when generating long reads for hybrid assembly.

Multiplexed ONT sequencing holds the promise of allowing complete and accurate genomes to be obtained from a single platform. Our results suggest that both *in silico* demultiplexing and laboratory kits need to improve before this is a reliable alternative to hybrid sequencing. Such development will be critical to ensuring the viability of ONT sequencing, particularly in routine clinical settings in the future. It has previously been hypothesised that the bimodal distribution observed in quality scores of reads (as calculated by their kmer identity to Illumina reads) delineates ‘good’ from ‘junk’ reads [9]. We speculate that in fact ONT reads with low identity to Illumina reads represent cross-barcode contamination. The long-read assembly problem is somewhat improved by consensus demultiplexing using two tools, but this is resource intensive, increases reads binned as ‘unclassified’ and is still not completely reliable. Hybrid assemblies are much less vulnerable to cross-barcode contamination which appears to be effectively removed by Unicycler’s process of mapping long reads to the short read assembly. Whilst reasonably high-quality long-read only assemblies can be achieved by running a single isolate per flowcell with subsequent polishing steps, the cost of this would currently be significantly higher than hybrid sequencing.

Different de-multiplexing, filtering and assembly parameters can produce different assemblies from the same input data. Whilst our assembly of the MGH757878 reference was very similar to the published sequence, further benchmarking of the effect of using different parameters is required but beyond the scope of this project. We included only a single *K. pneumoniae* reference strain meaning that the ground truth for most assemblies we performed was unknown, though notably in our first library overall structures did not change with an additional 24 hours of sequencing. An additional limitation is that we used a different extraction method for library one compared to all other libraries; however, the similar results obtained also demonstrate fully automated DNA extraction could be deployed to facilitate high-throughput hybrid sequencing workflows.

In conclusion we have demonstrated that high quality hybrid assemblies can be generated with much shorter sequencing times than are currently standard. The new ONT wash kit appears highly effective even to the point where reuse of the same barcodes on a flowcell seems acceptable when acquiring long reads for hybrid assemblies. Reusing flowcells for multiple libraries produces substantial potential per isolate cost reductions. Ultimately the opportunity to take advantage of this and conduct large-scale studies incorporating hybrid assembly is likely to help better inform future efforts to tackle some of the most important human pathogens.

## Authors Statements

### Authors and contributors

Conceptualization: SL, NSt, ASW; methodology: SL, NS; software: JS,NSa; formal analysis: SL; investigation: SL; resources: ASW, DC, TP, MA, MM; data curation: HP, KC, LB, DG, SL JK, MM; writing – original draft preparation: SL; writing – review and editing: All authors; supervision: DC, TP, ASW, NS; funding: SL, NS, ASW, DC, TP

## Data Bibliography

1. Raw sequencing data: NCBI BioProject Accession PRJNA604975
2. De novo assemblies: Figshare https://doi.org/10.6084/m9.figshare.11816532.v1

