## Supplement for "Optimised use of Oxford Nanopore Flowcells for Hybrid Assemblies"

### Supplementary methods

*Comparison of different demultiplexing/filtering options*

We compared four alternative filtering strategies across all five libraries:

- Guppy only: all raw reads as demultiplexed by guppy
    - Guppy + Deepbinner [1]: raw reads as demultiplexed by guppy with unclassified reads reassigned where possible by Deepbinner
    - Guppy + Deepbinner + Filtlong: as for option two but additionally processed with Filtlong using the following filters:
        - 90% of reads (based on highest quality score as judged by kmer match to Illumina reads) or maximum of 250Mbp (~ 50X depth), whichever resulted in fewer reads.
        - Only reads > 1000bp
        - –trim and –split 100
        - min_mean_q 25
    - Guppy + Deepbinner + random subsampling: as for option two but Rasusa[2] was used to subsample reads to ~ 30X coverage.

Using Guppy and Guppy+Deepbinner to reclaim unclassified reads provided the most complete assemblies (i.e. chromosome and all contigs circularised 36/57 and 37/57 respectively). Adding quality and length based filtering to Guppy + Deepbinner resulted in a slightly worse performance (33/57 assemblies circularised). Random sub-sampling to 150Mb with Rasusa provided the least complete assemblies (30/57) although in some cases the sampling threshold was more than the total number of sequenced bases for the sample. Using only the Guppy and Guppy+Deepbinner read filtering strategies we additionally compared assemblies created with Unicycler's --mode set to 'normal' and 'bold'. As expected, there were more complete assemblies using bold (36/57 (Guppy only) 37/57 (Guppy + Deepbinner)) compared with normal modes (29/57 (Guppy only) 31/57 Guppy + Deepbinner).

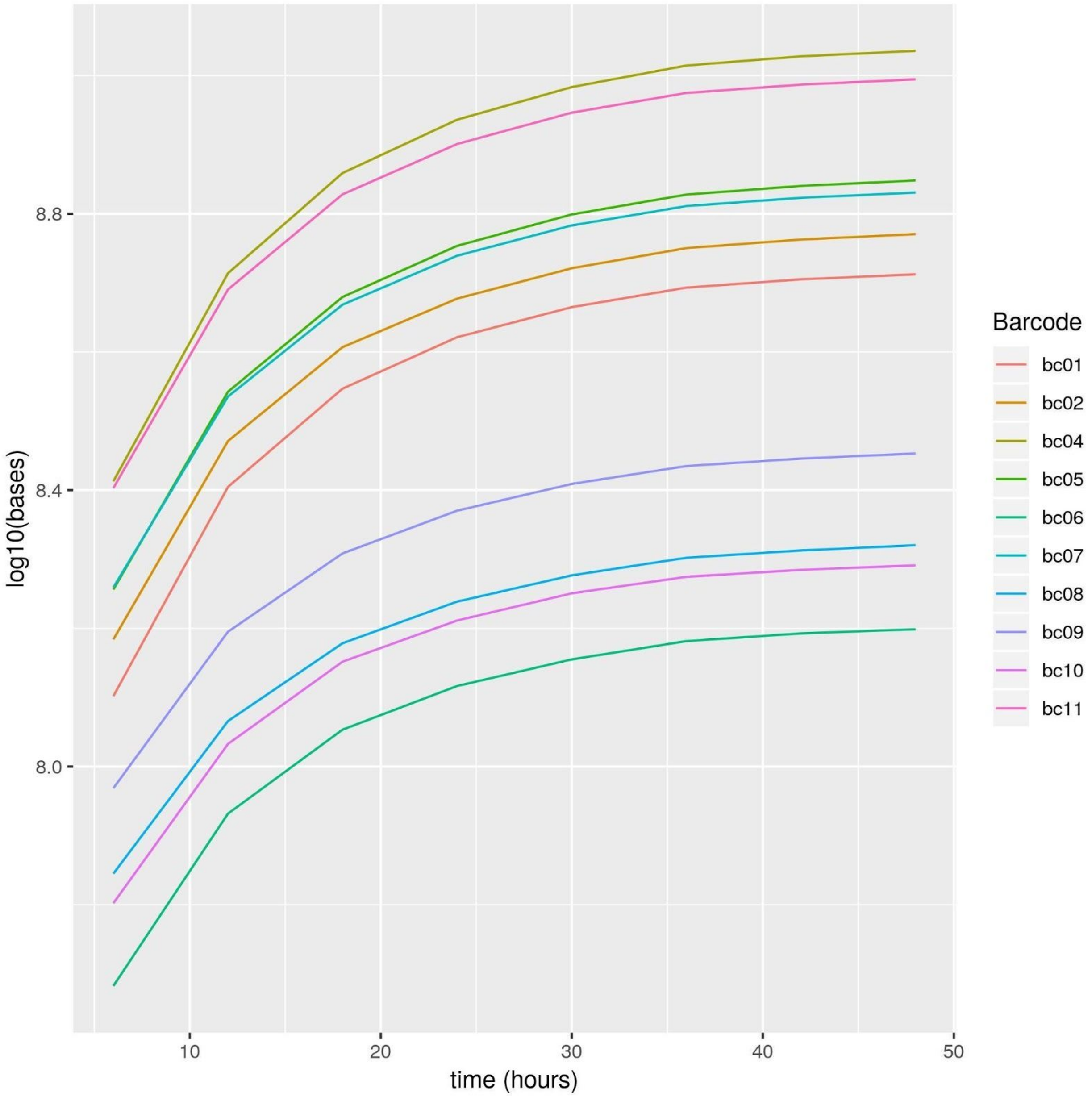

***Figure S1****: Log base output per barcode over time for library 1.*

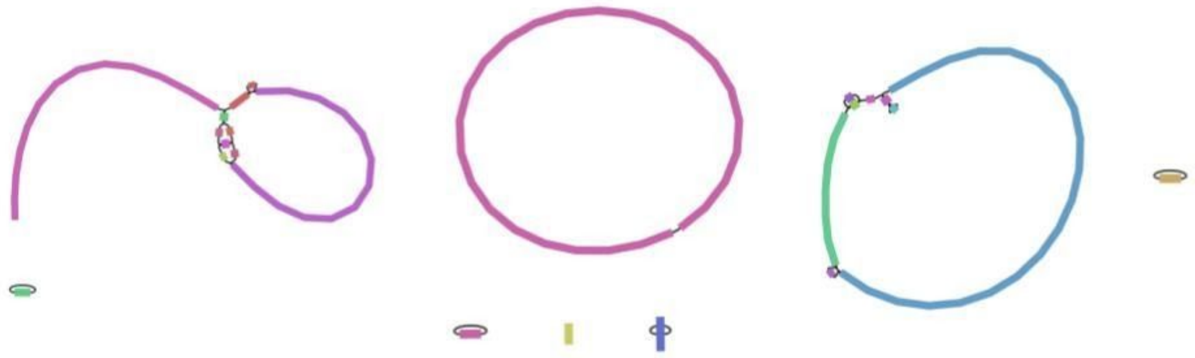

***Figure S2****: Graphs of the three incomplete assemblies from library 1 (left to right: barcodes bc02, bc04, bc05, isolates blc-23, blc-24, blc25) at 24 hours.*

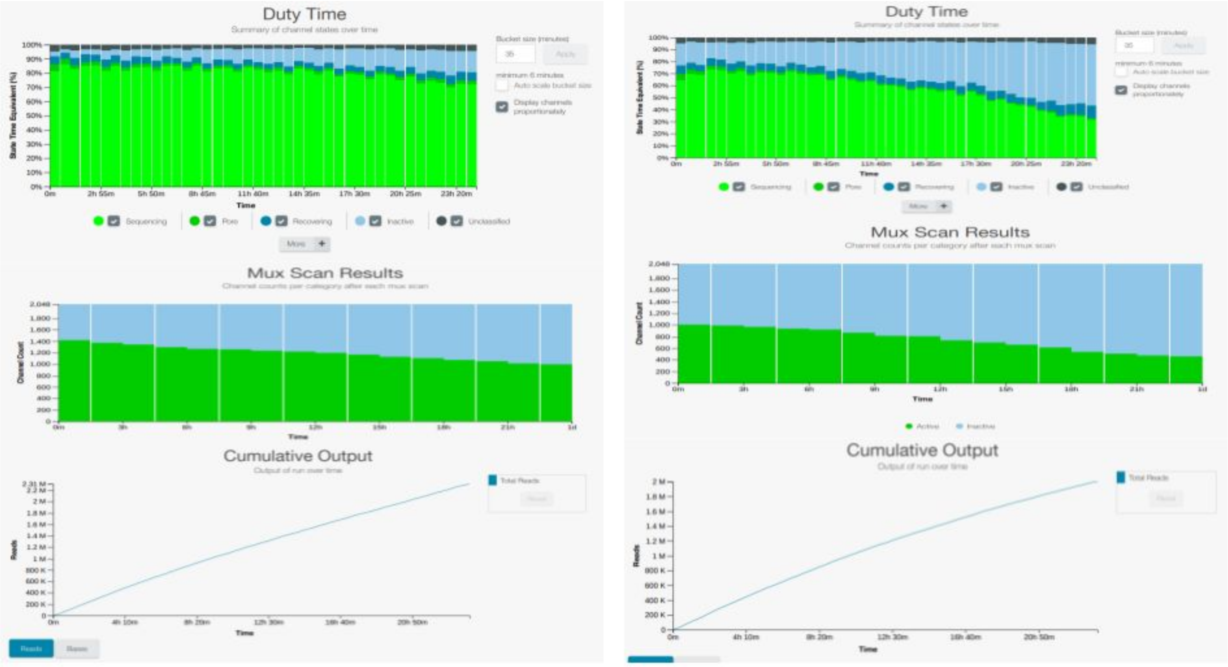

***Figure S3****: screenshots from the MinKnow reports from library 3 (left) and library 4 (right). These two libraries were run on the same flow cell with the ONT wash kit used following the end of library 3 sequencing at 24 hours before loading library 4.*

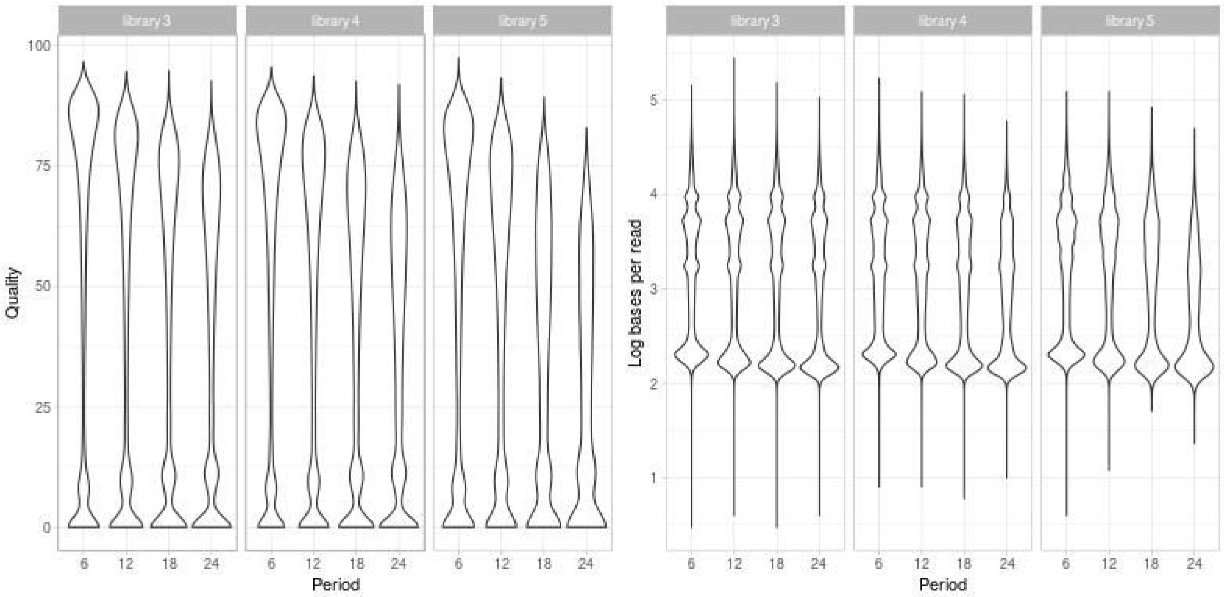

***Figure S4****: Read length/quality violin plots for libraries 3, 4 and 5 which were all sequenced consecutively on the same flowcell. Period 6 is time >= 0 hours < 6 hours, period 12 is time >=6 hours <12 hours etc.*

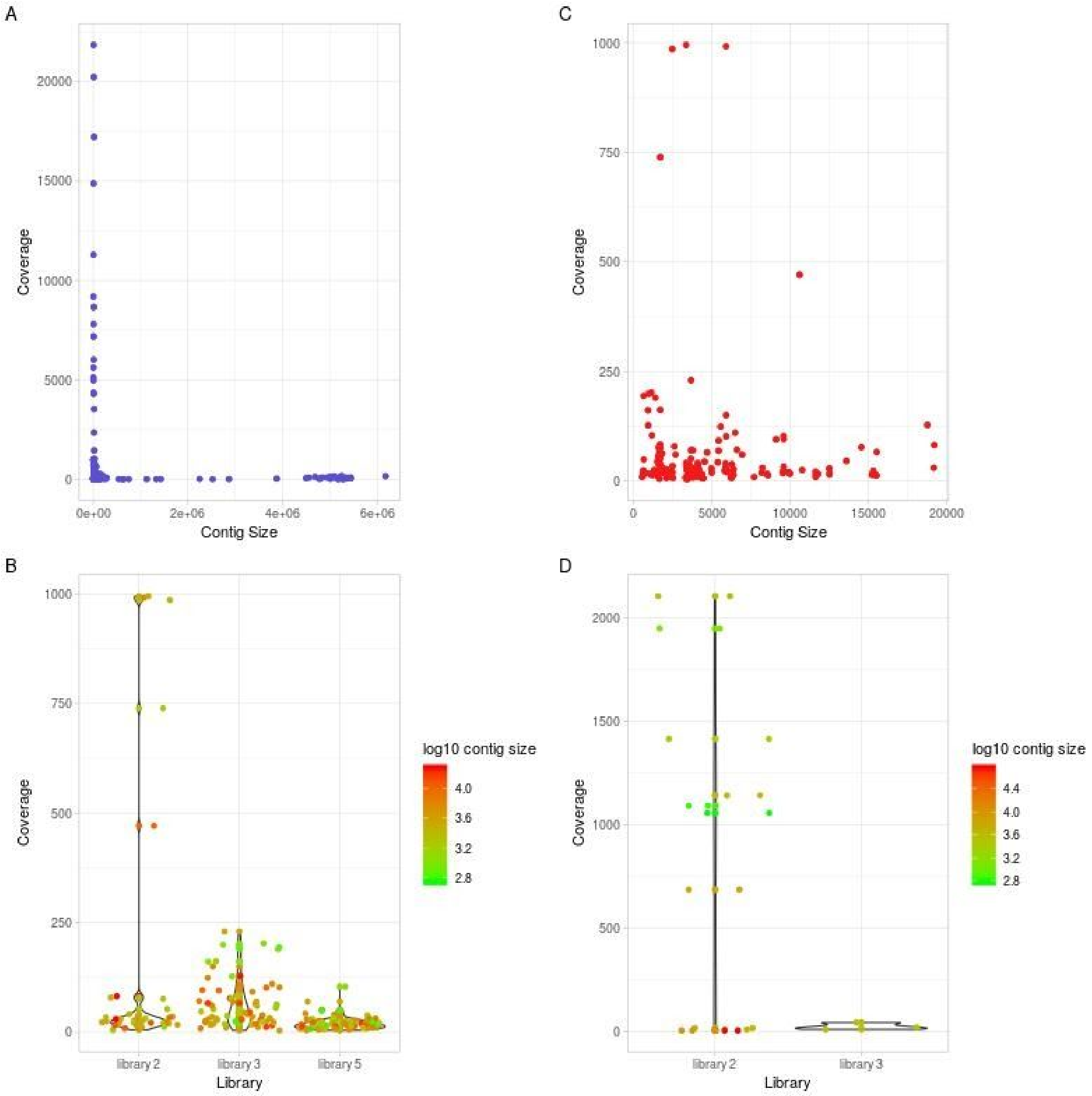

***Figure S5****: A - Contigs from Flye long-read only assemblies were classed as 'true' if they were matched to a contig in the hybrid assembly (match defined as an alignment of at least 100 kbp or ¼ of the replicon using the analysis.py script from [3]). Contigs were classed as unmapped if they had minimap reporting no mapping between them and the hybrid assembly. In general, 'true' small contigs were present at high coverage*

*compared to the spurious unmapped small contigs. B - Violin plot of the coverage of the unmapped contigs for each of the libraries (library 1 = 0, library 2 =36, library 3 = 67, library 5 = 78) using reads as demultiplexed by Guppy alone (there were no unmapped contigs for library 1). Log10 contig size is indicated by the colour gradient as shown. C - Violin plot of coverage of unmapped contigs using reads demultiplexed by taking the consensus of Deepbinner and Guppy. Flye assemblies from 1 and 5 produced no spurious contigs using this method and there were also substantially fewer for libraries 2 and 3 (n=22 and 3 respectively).*

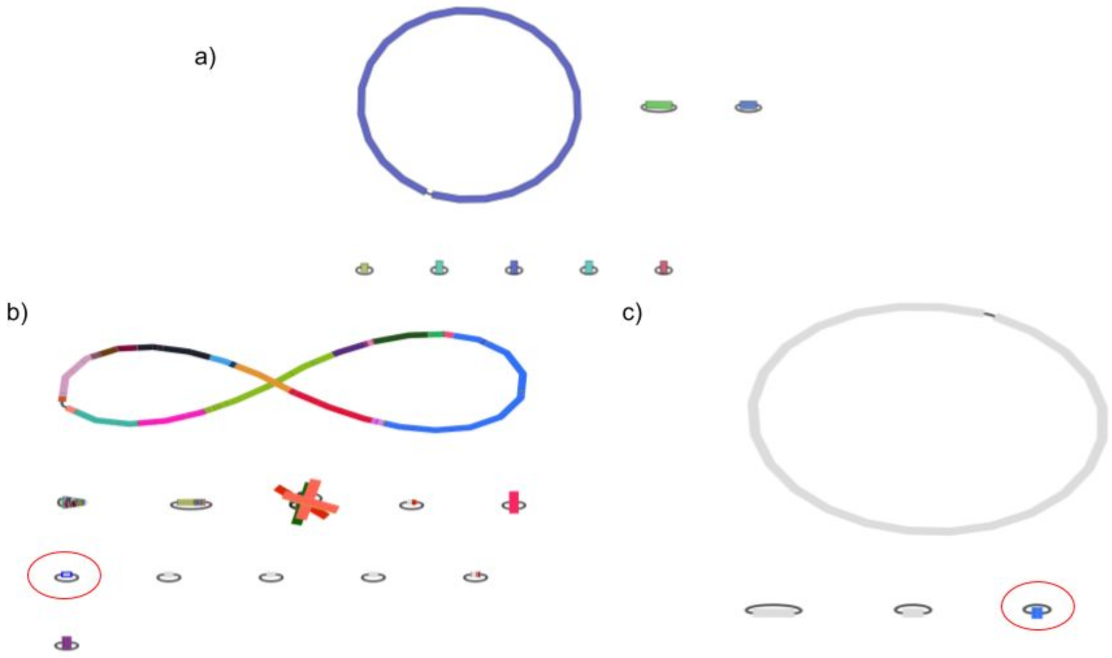

***Figure S6****: a) Hybrid assembly of blc-48 (barcode 02, library 3) showing the likely ground truth of a chromosome and 7 plasmids. b) Flye assembly of the same isolate. Colours represent blast hits from the short-read only assembly (i.e. replicons coloured grey like the one highlighted with a red ring are not present in the short read assembly and are presumed to be spurious). c) The contig highlighted with the red ring in b) was blasted against all other hybrid assemblies created from the same library (3), revealing a close match to a plasmid from blc-51 (barcode 05) and demonstrating likely cross contamination between barcodes in the same library. Note this does not represent between library contamination because library 3 was run on a new flow cell.*

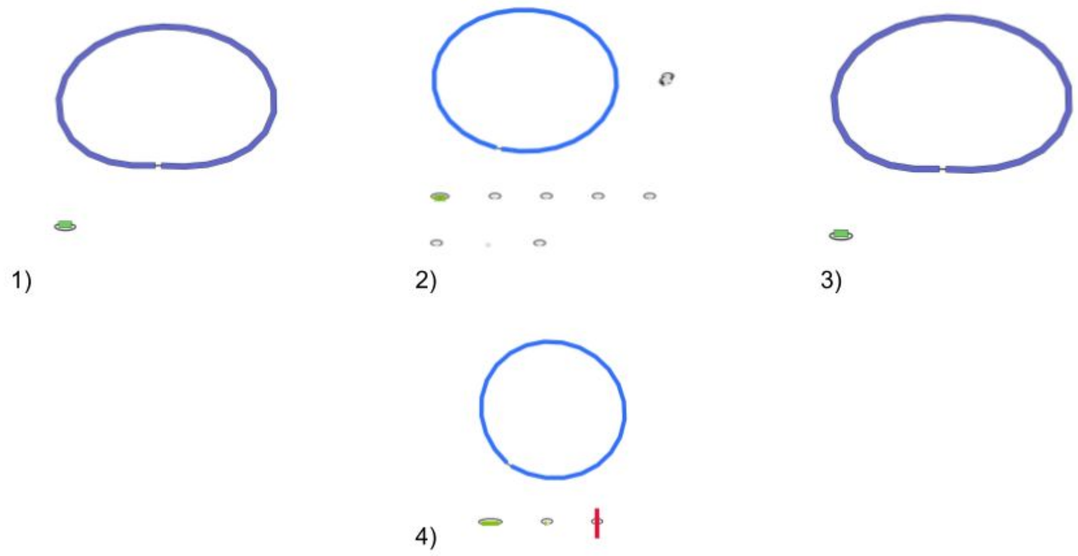

***Figure S7***: *1) Hybrid assembly of blc-50 (barcode04, library 3) representing the likely ground truth (1 chromosome, 1 plasmid). 2) Flye assembly of the same isolate - colours show blast hits using the against the hybrid assembly demonstrating the presence of the true chromosome and plasmid but also seven other likely spurious replicons. 3) Flye assembly using only reads assigned to barcode 04 by both Guppy and Deepbinner (1 chromosome, 1 plasmid). In general such a consensus demultiplexing strategy greatly improved long-read only assemblies. 4) blc-49 (barcode 03, library 3) using consensus demultiplexing of Guppy and Deepbinner - colours represent blast hits against the short-read only assembly. There is a spurious additional part of which is green (ie truly belongs to the plasmid on the left) and part is grey (ie is not represented in the hybrid assembly). This demonstrates that whilst consensus demultiplexing is useful to improve*

*the accuracy of multiplexed long read only assembly, it is likely still prone to within-library contamination.*

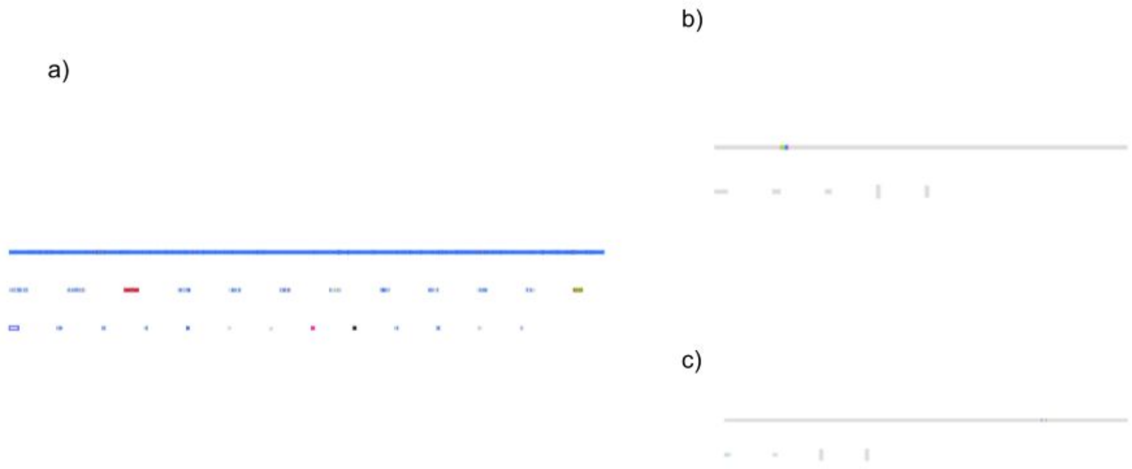

***Figure S8****: a) Flye assembly of RBHSTW-0000059 from the REHAB dataset [4] (multiplexed with ONT's native barcoding kit). Replicons appear linear because a fasta file (downloaded from the study's figshare page (https://figshare.com/articles/Hybrid_Enterobacteriaceae_assemblies_using_PacBio_Illumina_or_ONT_Illumina_sequencing/7649051) was loaded into bandage rather than a gfa file. Colours represent blast hits against the subsampled Nanopore/Illumina hybrid assembly. Replicons which appear grey are not represented in the hybrid assembly. b) the contig highlighted in a blue box in a blasted against the MGH78578 hybrid assembly, demonstrating a near perfect hit to this isolates chromosome. c) The same*

*contig blasted against the ONT/Illumina subsampled hybrid assembly of RBHSTW-0000059 demonstrating only very small and non-continuous hits representing false positive matches. This demonstrates that the problem of within library contamination affecting long read only assemblies is not unique to the rapid barcoding kit nor to this study.*

| Library | Isolate Name | Barcode | Species | MLST | Accession Number |
|---|---|---|---|---|---|
| 1 | blc-22 | barcode01 | *K. pneumoniae* | 38 | |
| 1 | blc-23 | barcode02 | *E. coli* | 73 | |
| 1 | blc-24 | barcode04 | *K. oxytoca* | - | |
| 1 | blc-25 | barcode05 | *E. coli* | 5430 | |
| 1 | blc-26 | barcode06 | *E. coli* | 4219 | |
| 1 | blc-27 | barcode07 | *Enterobacter cloacae* | 422 | |
| 1 | blc-28 | barcode08 | *Proteus spp.* | - | |
| 1 | blc-29 | barcode09 | *Aeromonas veronii* | - | |
| 1 | blc-30 | barcode10 | *E. coli* | 73 | |
| 1 | blc-31 | barcode11 | *K. pneumoniae* | - | |
| 2 | blc-32 | barcode01 | *K. pneumoniae* | 490 | |
| 2 | blc-33 | barcode02 | *E. coli* | 80 | |
| 2 | blc-34 | barcode03 | *K. pneumoniae* | 17 | |
| 2 | blc-35 | barcode04 | *K. pneumoniae* | 490 | |
| 2 | blc-36 | barcode05 | *K. pneumoniae* | 490 | |
| 2 | blc-37 | barcode06 | *K. pneumoniae* | 490 | |
| 2 | blc-38 | barcode07 | *K. pneumoniae* | 490 | |
| 2 | blc-39 | barcode08 | *E. coli* | 69 | |
| 2 | blc-40 | barcode09 | *K. pneumoniae* | 490 | |
| 2 | blc-41 | barcode10 | *E. coli* | 127 | |
| 2 | blc-42 | barcode11 | *E. coli* | 95 | |
| 2 | blc-43 | barcode12 | *K. pneumoniae* | 490 | |
| 3 | blc-53 | barcode07 | *E. coli* | 127 | |
| 3 | blc-54 | barcode08 | *E. coli* | 88 | |
| 3 | blc-55 | barcode09 | *K. pneumoniae* | 490 | |
| 3 | blc-44 | barcode10 | *E. coli* | 420 | |
| 3 | blc-45 | barcode11 | *K. pneumoniae* | 490 | |

| | | | | |
|---|---|---|---|---|
| 3 | blc-46 | barcode12 | *E. coli* | 372 |
| 3 | blc-47 | barcode01 | *K. pneumoniae* | 490 |
| 3 | blc-48 | barcode02 | *K. pneumoniae* | 490 |
| 3 | blc-49 | barcode03 | *K. pneumoniae* | 45 |
| 3 | blc-50 | barcode04 | *E. coli* | 127 |
| 3 | blc-51 | barcode05 | *K. pneumoniae* | 15 |
| 3 | blc-52 | barcode06 | *E. coli* | 428 |
| 4 | blc-44 | barcode01 | *E. coli* | 420 |
| 4 | blc-45 | barcode02 | *K. pneumoniae* | 490 |
| 4 | blc-46 | barcode03 | *E. coli* | 372 |
| 4 | blc-47 | barcode04 | *K. pneumoniae* | 490 |
| 4 | blc-48 | barcode05 | *K. pneumoniae* | 490 |
| 4 | blc-49 | barcode06 | *K. pneumoniae* | 45 |
| 4 | blc-50 | barcode07 | *E. coli* | 127 |
| 4 | blc-51 | barcode08 | *K. pneumoniae* | 15 |
| 4 | blc-52 | barcode09 | *E. coli* | 428 |
| 4 | blc-53 | barcode10 | *E. coli* | 127 |
| 4 | blc-54 | barcode11 | *E. coli* | 88 |
| 4 | blc-55 | barcode12 | *K. pneumoniae* | 490 |
| 5 | blc-56 | barcode01 | *K. pneumoniae* | 490 |
| 5 | blc-57 | barcode02 | *E. coli* | 127 |
| 5 | blc-58 | barcode03 | *K. pneumoniae* | 15 |
| 5 | blc-59 | barcode04 | *K. pneumoniae* | 45 |
| 5 | blc-60 | barcode05 | *K. pneumoniae* | 490 |
| 5 | blc-61 | barcode06 | *K. pneumoniae* | 490 |
| 5 | blc-62 | barcode07 | *E. coli* | 420 |
| 5 | blc-63 | barcode08 | *E. coli* | 127 |
| 5 | blc-64 | barcode09 | *E. coli* | 372 |
| 5 | blc-65 | barcode10 | *E. coli* | 428 |
| 5 | blc-66 | barcode11 | *E. coli* | 88 |
| 5 | blc-67 | barcode12 | *K. pneumoniae* | 490 |

***Table S1****: Summary of isolates sequenced in each library. * reference strain MGH75878*

| Library | Human reads | Total Reads | % human reads |
|---|---|---|---|
| 1 | 553 | 1648038 | 0.02 |
| 2 | 147 | 818091 | 0.02 |
| 3 | 357 | 1586627 | 0.02 |
| 4 | 241 | 1328083 | 0.02 |
| 5 | 62 | 331211 | 0.02 |

***Table S2**: Reads binned as human by centrifuge from each library. Library 2 was run on a flow cell first used to sequence a human pathology specimen for 24 hours which had 2028024/2059966 (98.4%) initial reads binned as human. After washing the reads classified as human appeared within the range of all other libraries (which were sequenced on flow cells on which no human DNA had been loaded). This demonstrates the highly effective removal of DNA by the ONT wash kit.*
